# The farnesyl transferase inhibitor KO-2806 re-sensitizes relapsing tumors to RAS inhibition

**DOI:** 10.1101/2024.12.20.629824

**Authors:** Hetika Vora Patel, Alison Elizabeth Smith, Stacia Chan, Jovylyn Gatchalian Gasendo, Tejas Samantaray, Linda Kessler, Amitava Mitra, Xuefeng Zhu, Yahu A. Liu, Francis Burrows, Shivani Malik

## Abstract

Resistance remains a key issue limiting the clinical benefit from RAS-targeting therapeutic agents and necessitates combination approaches. We identify persistent mTORC1 activity in preclinical *KRAS*-mutant NSCLC and CRC models as a frequent, nongenetic driver of inherent and adaptive resistance to RAS inhibition. This vulnerability is targetable with the farnesyl transferase inhibitor KO-2806, which blocks mTORC1 activation via RHEB while sparing mTORC2 and its associated toxicities. The addition of KO-2806 to NSCLC or CRC tumors progressing on mutant-selective RAS inhibitors led to rapid and durable tumor regression. In contrast, switching from mutant-selective to pan-RAS inhibitor monotherapy resulted in only stasis of NSCLC tumors and had no effect on CRC tumor progression. Further, the addition of KO-2806 rescued sensitivity of progressing tumors to the pan-RAS inhibitor RMC-6236. Our results establish mTORC1 as an important mediator of escape from RAS inhibition and highlight KO-2806 as a promising RAS companion inhibitor in patients with prior RAS inhibitor exposure.

**Significance:** Utilizing *in vivo* models of tumor relapse, we define a subset of RAS inhibitor-resistant tumors in which vertical inhibition of MAPK is insufficient to restore sensitivity. By controlling parallel mTORC1 activity, KO-2806 may expand utility of RAS inhibitors in patients that have progressed on RAS-targeted therapy, regardless of inhibitor class.

## INTRODUCTION

The approval of KRAS^G12C^ inhibitors and ongoing clinical development of additional mutant-selective and pan-RAS targeted agents has heralded a shift in the treatment paradigm for patients with RAS-mutant tumors(1–4). Nevertheless, only a subset of these patients respond to RAS inhibitor therapy, and most responses are limited in duratione(5). This limitation has led to a flurry of proposed combination strategies to mitigate resistance, many of which center around targeting nodes upstream or downstream of oncogenic KRAS to inhibit pathway output more potently(5–7).

In preclinical models of non-small cell lung cancer (NSCLC), it has been demonstrated that the degree of inhibition of mTORC1 correlates with responsiveness to KRAS^G12C^-targeted agents, and inhibitors of KRAS^G12C^ and mTOR are synergistic(8). Moreover, mutations that co-occur with *KRAS*, like *STK11* in NSCLC and *PIK3CA* in colorectal cancer (CRC), are thought to limit KRAS^G12C^ inhibitor efficacy, pointing to the importance of mTOR in therapy response and resistance in these tumor types(9, 10). In CRC specifically, elevated basal receptor tyrosine kinase (RTK) activity reduces sensitivity to KRAS inhibition by swiftly reactivating downstream signaling(11–13). Although pan-RAS inhibitors should restrict activation of wildtype RAS previously implicated in KRAS^G12C^ inhibitor resistance, persistent mTORC1 signaling likely remains a liability regardless of inhibitor mechanism(4, 13).

We have previously shown that farnesyl transferase inhibition enhances the efficacy of PI3Kα inhibitors by blocking RHEB-mediated activation of mTORC1(14). We speculate that the farnesyl transferase inhibitor KO-2806 (currently under clinical investigation in a phase 1 trial to assess monotherapy and combination therapies in solid tumors, NCT06026410) may similarly improve response to RAS inhibitors in *KRAS*-mutant NSCLC and CRC given the demonstrated role of mTORC1 in these indications. However, as more patients are treated with KRAS^G12C^- specific inhibitors and additional RAS-targeted agents gain regulatory approval and become part of the treatment landscape, the need for viable therapeutic options after failure on a (K)RAS inhibitor will expand. Here, we ask whether farnesyl transferase inhibition can re-sensitize relapsing *KRAS*-mutant NSCLC and CRC tumors to multiple classes of RAS inhibitors.

## RESULTS

### KO-2806 is a potent, selective farnesyl transferase inhibitor with potential to enhance KRAS inhibitor efficacy

The structure of the farnesyl transferase inhibitor (FTI) KO-2806 is shown in Fig. 1A. We first assessed the compound’s activity in biochemical and cellular assays. KO-2806 potently inhibited *in vitro* farnesyl transferase (FTase) activity (IC_50_ 2.4 nM), while sparing activity of geranylgeranyl transferase (GGTase), a homologous prenyl transferase (IC_50_ >1000 nM), indicating marked selectivity (Fig. 1B). As HRAS is a *bona fide* farnesylation-dependent oncogenic driver, we exposed the HRAS-mutant bladder carcinoma cell line T24/83 to increasing doses of KO-2806 *in vitro*. Both cell viability (Fig. 1C) and farnesylation of key targets (indicated by mobility shift of HRAS and RHEB or appearance of unprocessed prelamin A(15), Fig. 1D) were inhibited in a dose-dependent manner. *In vivo*, KO-2806 also blocked the growth of HRAS-mutant head and neck patient-derived xenograft (PDX) tumors, further pointing to on-target activity (Fig. S1).

**Figure 1.**
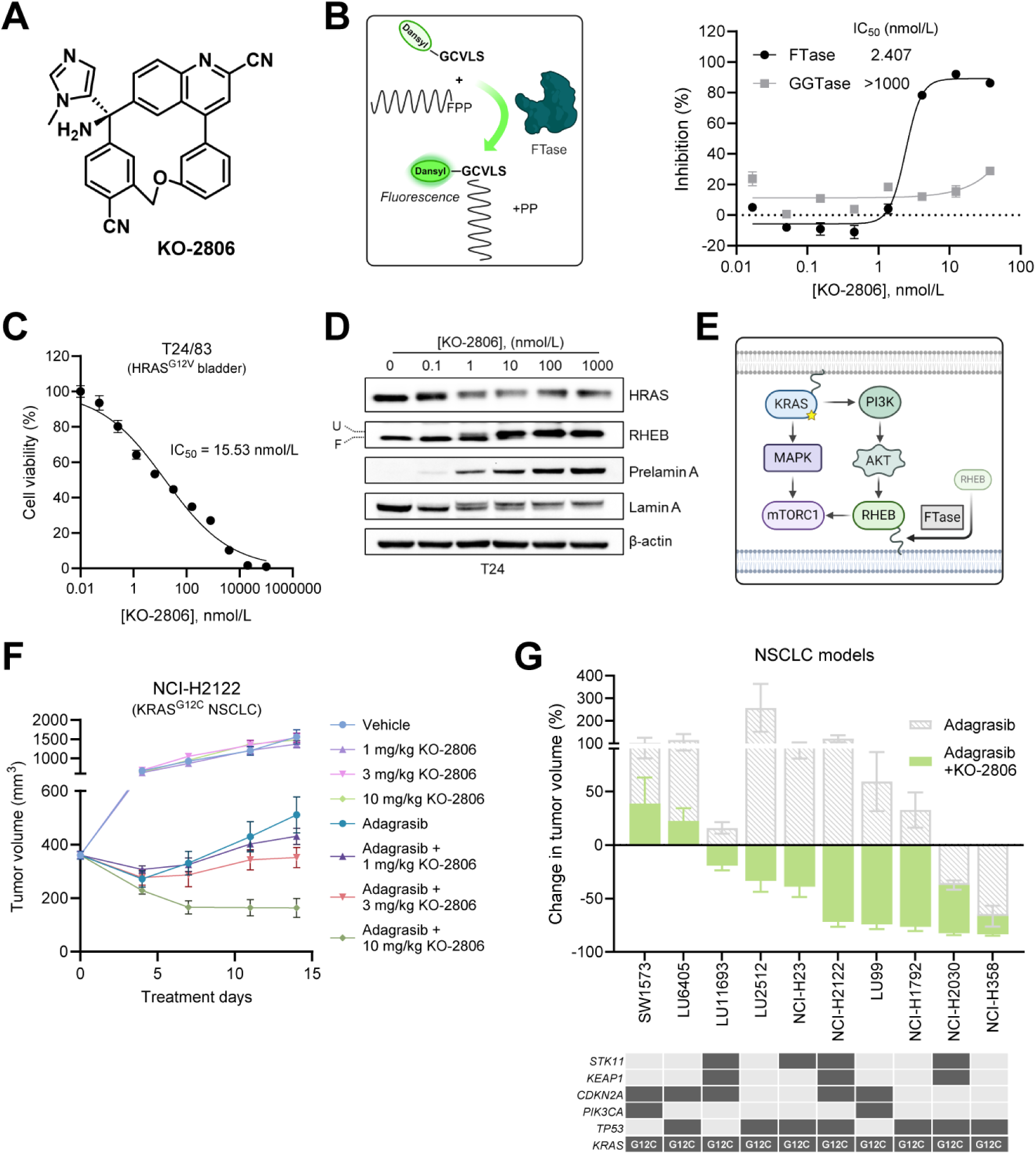
KO-2806 is a potent farnesyl transferase inhibitor that deepens responses of NSCLC models to KRAS^G12C^ inhibition. **A**, Chemical structure of KO-2806. **B**, Graphical overview and results of farnesyl transferase (FTase) biochemical activity fluorescence assay using farnesyl pyrophosphate (FPP) and dansyl-labeled peptide substrates. Increasing concentrations of KO-2806 were added to FTase and geranylgeranyl transferase (GGTase) activity assays. Nonlinear regression with log(inhibitor) vs. response was used to calculate IC_50_. Data are means ± SEM of 6-10 replicates. **C**, Inhibition of proliferation of T24/83 cells treated with increasing doses of KO-2806 for 4 days. Data are means ± SEM of 10 biological replicates. **D**, Immunoblot of the indicated farnesylated proteins and loading control in T24/83 cells treated with increasing concentrations of KO-2806 for 48 hours. Mobility shift of unfarnesylated (U) vs. farnesylated (F) RHEB indicated to left of image. Images are representative of 2 biological replicates. **E**, Graphical summary of KRAS-mTORC1 signaling, including farnesylated RHEB’s role in activating mTORC1. **F**, Growth of NCI-H2122 xenograft tumors treated with the indicated doses of KO-2806 (PO BID) as monotherapy or in combination with 100 mg/kg adagrasib (PO QD). Data are mean tumor volumes ± SEM, *n* = 6 mice per group. **G**, Tumor efficacy waterfall plots of *KRAS^G12C^*-mutant NSCLC xenograft models treated with 30-100 mg/kg adagrasib as monotherapy (gray bars) or in combination with 10-15 mg/kg KO-2806 (colored bars). Each bar represents the mean percent change in tumor volume at day 25-35 relative to baseline (day 0), ± SEM, *n* = 6-8 animals per group. Oncoplots illustrating the mutational status of key driver genes in each model are shown below each waterfall plot. Mutations are indicated by dark gray shading.

The FTI tipifarnib was shown to be active as a monotherapy in patients with HRAS-mutant cancers(16), but our prior work suggests the broader utility of FTIs lies in combination with targeted agents and hinges upon the ability of the FTIs to inhibit mTORC1 via perturbation of RHEB localization(14) (Fig. 1E). Given the documented role mTORC1 plays in mediating sensitivity to KRAS^G12C^ inhibitors(8, 17), we hypothesized that KO-2806 may help overcome resistance to these inhibitors. To address this possibility, we began by treating the *KRAS^G12C^*- mutant NSCLC xenograft model NCI-H2122 with the FDA-approved G12C-selective inhibitor adagrasib, KO-2806, or the combination (Fig. 1F). While single agent adagrasib (100 mg/kg) initially decreased tumor volume, tumors began to regrow within one week. The addition of KO- 2806 dose-dependently slowed this regrowth, and a twice daily 10 mg/kg dose in combination with adagrasib resulted in deep tumor regressions. We further explored the potential of this combination by administering KO-2806 and adagrasib to a panel of *KRAS^G12C^*-mutant NSCLC xenograft models with varying sensitivity to adagrasib monotherapy as well as a spectrum of co-mutations (Fig. 1G). In all models tested, KO-2806 enhanced tumor growth inhibition or deepened regressions observed with single agent adagrasib. The combination was generally well tolerated (Fig. S2). The breadth and magnitude of this effect supports clinical investigation of KO-2806 as a combination partner for KRAS^G12C^-selective inhibitors.

### Compensatory mTOR signaling is robustly inhibited by the combination of KO-2806 and adagrasib in *KRAS^G12C^* NSCLC models

We next probed the molecular mechanism underlying the enhanced antitumor effect of combined KO-2806 and adagrasib using *in vitro* and *in vivo* systems. In *KRAS^G12C^* NSCLC cells cultured as tumor spheroids, adagrasib potently and durably inhibited RAS (phosphorylation of ERK and p90 RSK) activity for 48 hours (Fig. 2A, S3A). In contrast, inhibition of mTORC1 substrate phosphorylation (p70 S6K, 4E-BP1, and S6) was incomplete and/or transient. KO- 2806 exposure reduced basal phosphorylation of these mTOR substrates and augmented both the depth and duration of inhibition by adagrasib. Greater mTOR inhibition by the adagrasib-KO- 2806 combination corresponded to a loss of RHEB farnesylation (as indicated by mobility shift), induction of apoptosis (caspase and PARP cleavage), and cell cycle arrest (decrease in cyclin D1 levels and Rb phosphorylation), suggesting KO-2806 enhances KRAS^G12C^ inhibitor activity by restricting RHEB-dependent mTORC1 activation. To explore this, we asked whether genetic depletion of *RHEB* expression in NCI-H2122 tumor spheroids would phenocopy the effects of KO-2806 on cell proliferation and signaling. Indeed, combined exposure to siRNAs against *RHEB* and adagrasib had similar inhibitory effects on proliferation (Fig. 2B) and mTOR substrate phosphorylation (Fig. 2C) as the combination of KO-2806 and adagrasib. Further, a farnesylation-deficient RHEB^C181S^ mutant (expressed in NCI-H2122 cells depleted of wildtype RHEB) exhibited a similar cellular localization (Fig. S3B), gel shift, and effect on mTOR signaling (Fig. S3C) as wildtype RHEB upon KO-2806 exposure. This indicates that loss of RHEB farnesylation due to inhibition of FTase activity likely mediates the observed phenotype in KO-2806-exposed *KRAS*-mutant cells.

**Figure 2.**
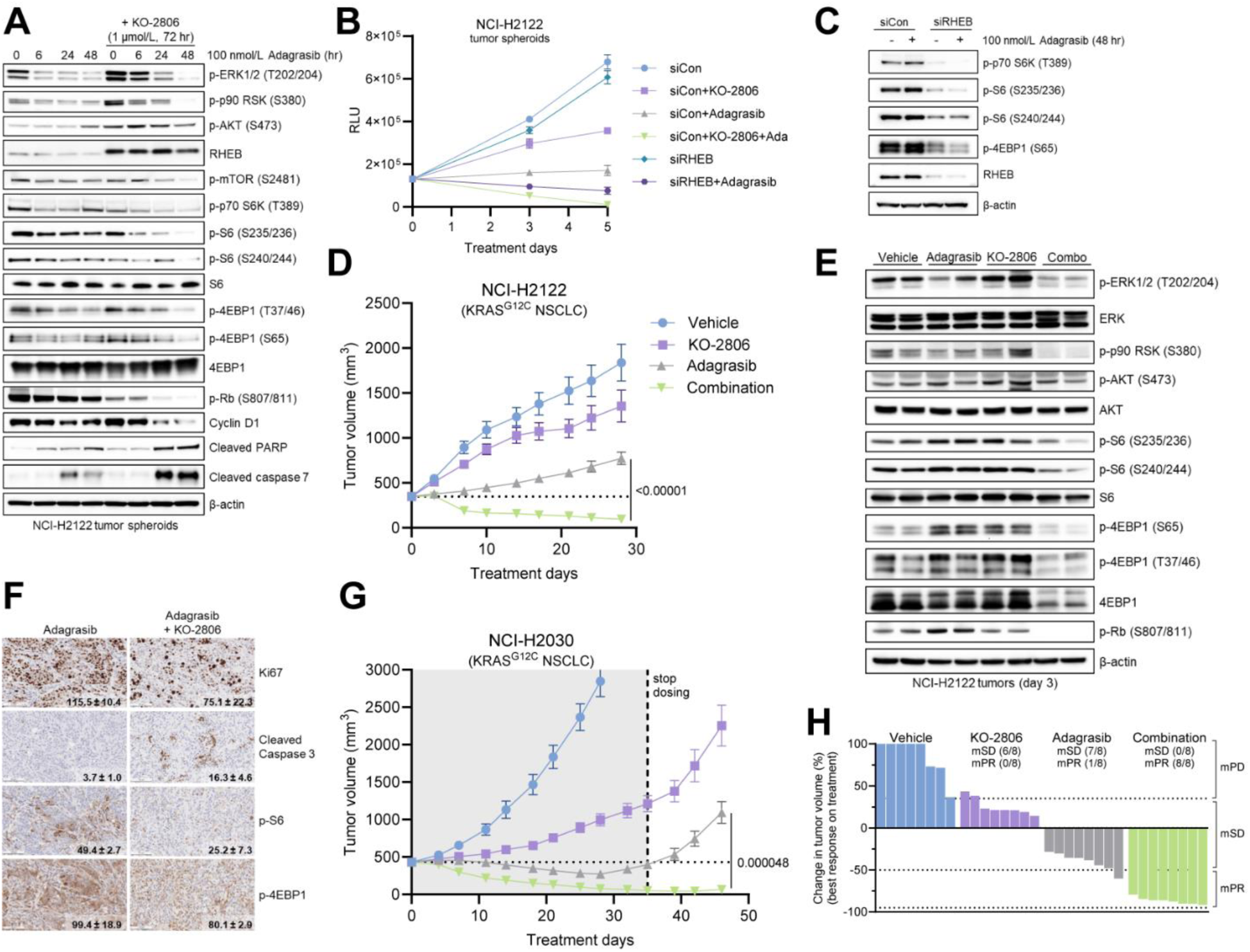
Combination of KO-2806 with adagrasib blunts compensatory mTOR signaling and induces apoptosis and regression in *KRAS^G12C^* NSCLC preclinical models. **A**, Immunoblots of the indicated signaling proteins in NCI-H2122 tumor spheroids exposed to 100 nmol/L adagrasib for 0, 6, 24, or 48 hours in the presence or absence of 1 μmol/L KO-2806. Images are representative of 3 biological replicates. **B**, Growth of NCI-H2122 spheroids transfected with siRNAs against *RHEB* (or a control nontargeting pool, siCon) in the presence or absence of 100 nmol/L adagrasib and 100 nmol/L KO-2806. Data are means ± SD of 6 biological replicates. **C**, Immunoblots of the indicated mTOR signaling proteins in cells from **B** in the presence or absence 100 nmol/L adagrasib for 48 hours. Images are representative of 3 biological replicates. **D**, Growth of NCI-H2122 cell-line derived xenograft tumors treated with 100 mg/kg adagrasib PO QD, 10 mg/kg KO-2806 PO BID, or the combination for 28 days. Each point represents mean tumor volume ± SEM, *n* = 8 mice per group. **E**, Immunoblots of the indicated MAPK and mTOR signaling proteins in NCI-H2122 tumors treated as specified in **D** for 3 days. Images are representative of 2 biological replicates. **F**, Immunohistochemistry analysis of NCI-H2122 tumors treated with adagrasib ± KO-2806 for 3 days. Quantification of staining is overlaid on representative images (mean H-scores ± SD, *n* = 3 tumors). Scale bar, 100 μm. **G**, Growth of NCI-H2030 xenograft tumors treated as in **D** for 35 days, followed by off-dosing observation to assess durability of treatment. Each point represents mean tumor volume ± SEM, *n* = 8 mice per group. Statistical comparison by unpaired Student’s t-test, p-values as indicated on plots. **H**, Waterfall plot of best responses (% change in tumor volume from baseline) of each animal on treatment in **G**. Response calls were made using mRECIST criteria and are indicated above the plot.

To validate this mechanism *in vivo*, we treated NCI-H2122 xenograft tumors with adagrasib, KO- 2806, or the combination (Fig. 2D, S3D). Although adagrasib monotherapy was only able to slow tumor growth, the combination of adagrasib and KO-2806 led to rapid tumor regression, which was maintained over the course of the study. Tumors were collected after three days of treatment to assess tumor cell signaling events that may have led to this regression. As was observed *in vitro*, mTOR activity (phospho-S6, 4E-BP1) was lower in tumors treated with combined KO-2806 and adagrasib than those treated with adagrasib alone, which correlated with diminished tumor cell proliferation (phospho-Rb, Ki67) and induction of apoptosis (caspase cleavage) (Fig. 2E and F). Finally, we identified a *KRAS^G12C^*-mutant xenograft model that exhibited a robust initial response to adagrasib monotherapy to ask whether KO-2806 could improve the durability of responses. NCI-H2030 xenograft tumors treated with adagrasib monotherapy regressed for several weeks but began to regrow at approximately day 28 (Fig. 2G and H, S3E). The addition of KO-2806 to adagrasib not only deepened tumor regression but also prevented this regrowth. The durability of these responses was further probed by suspending treatment after day 35. Adagrasib-treated tumors rapidly doubled in volume, while combination-treated tumors did not regrow. These results demonstrate that KO-2806 can deepen and extend responses to KRAS^G12C^ inhibition by disrupting RHEB-mediated mTORC1 activation.

### KO-2806 blocks rebound of mTOR signaling and deepens antitumor responses to mutant-selective and pan-RAS inhibitors in CRC

*KRAS^G12C^* mutations are rare outside of NSCLC(7). Thus, mutant-selective agents targeting more prevalent mutations, such as KRAS^G12D^, or pan-RAS/pan-KRAS inhibitors that block activity of mutant and wildtype RAS/KRAS may treat a more diverse set of patients in other *KRAS*-driven indications, such as CRC(12). We speculated that the synergy between KO-2806 and adagrasib we observed in NSCLC models might extend to RAS inhibitors of other classes. To begin, we assessed whether KO-2806 could potentiate the antitumor efficacy of a KRAS^G12D^- specific inhibitor, MRTX1133, or a pan-RAS(ON) inhibitor, RMC-6236, in models of *KRAS-* mutant CRC. Preclinical studies have linked the incomplete inhibition of S6 phosphorylation downstream of mTOR with diminished responses to these agents. As KO-2806 and a KRAS^G12C^-selective inhibitor potently inhibited mTOR activity in NSCLC cells *in vitro*, we first examined the dynamics of signaling inhibition by MRTX1133 in a G12D-mutant CRC cell line, GP2D (Fig. 3A). MRTX1133 inhibited phosphorylation of ERK for 48 hours, but inhibition of mTORC1 (phospho-p70 S6K, phospho-4EBP1, phospho-S6) was transient, observed at 4 hours before rebounding fully by 48 hours. Simultaneous exposure to MRTX1133 and KO-2806 resulted in deeper inhibition of mTOR substrates, and fully blocked rebound, correlating with cell cycle arrest (decreased phospho-Rb) and apoptosis (PARP cleavage). Similar trends were observed with pan-RAS inhibition; RMC-6236 was unable to durably inhibit mTOR activity, but the addition of KO-2806 controlled mTOR rebound (Fig. 3B). This effect was not limited to cells harboring *KRAS^G12D^* mutations and was also seen in *KRAS^G13D^*-mutant cells (Fig. S4A and B).

**Figure 3.**
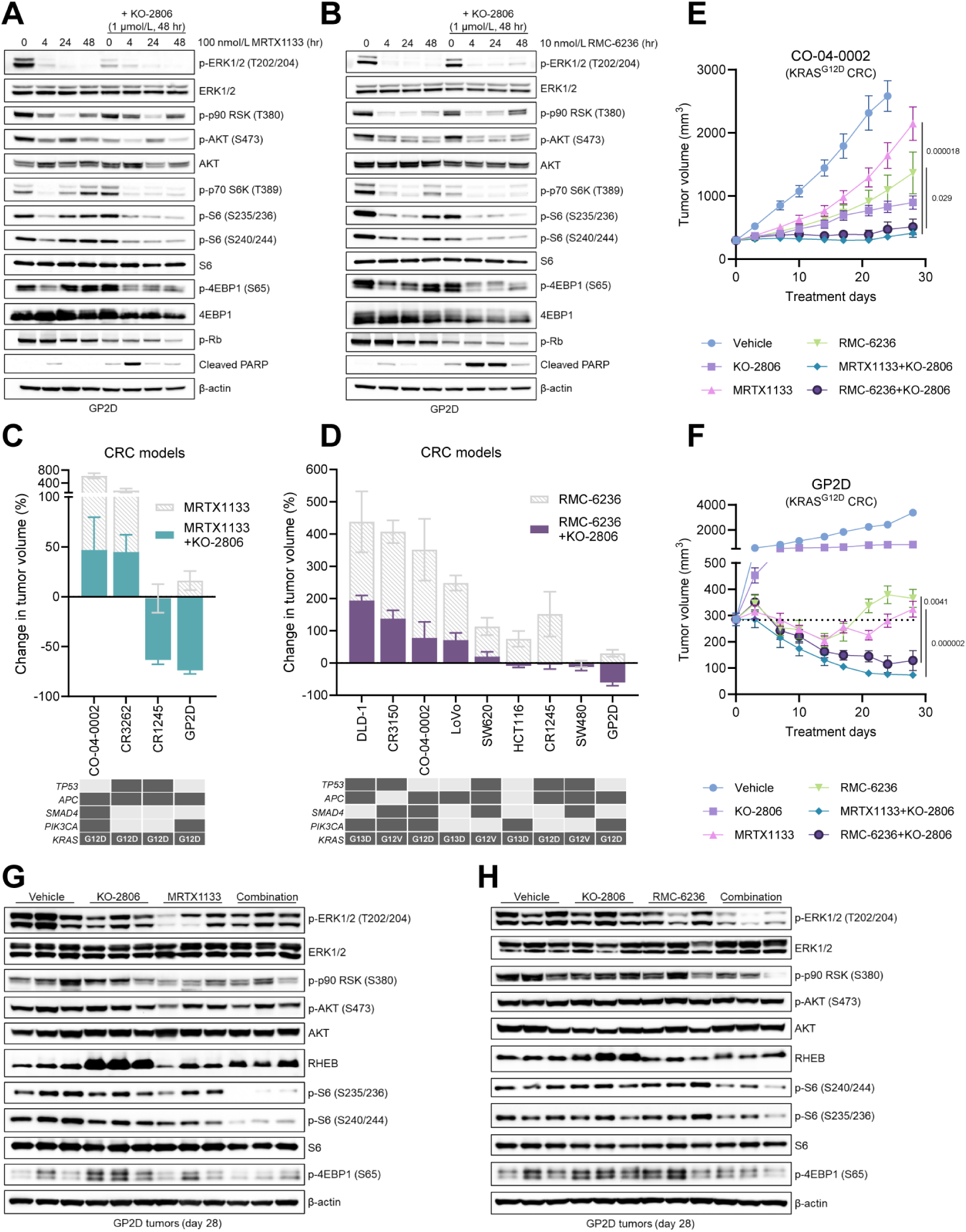
Farnesyl transferase inhibition blocks rebound of mTOR signaling following mutant-selective or pan-RAS inhibition in preclinical models of CRC, leading to enhanced tumor growth inhibition. **A** and **B,** Immunoblots of the indicated signaling proteins in GP2D CRC cells exposed to (**A**) 100 nmol/L MRTX1133 or (**B**) 10 nmol/L RMC-6236 for 0, 4, 24, or 48 hours in the presence or absence of 1 μmol/L KO-2806. Images are representative of 3 biological replicates. **C** and **D**, Tumor efficacy waterfall plots of *KRAS*-mutant CRC xenograft models treated with 10-30 mg/kg MRTX1133 IP BID (**C**) or 10-25 mg/kg RMC-6236 PO QD (**D**) as monotherapy (gray bars) or in combination with 10 mg/kg KO-2806 PO BID (colored bars). Each bar represents the mean percent change in tumor volume at day 28-35 (endpoint) relative to day 0, ± SEM, *n* = 8 mice per treatment group. Oncoplots illustrating the mutational status of key driver genes as well as the *KRAS* mutation codon in each model are shown below each waterfall plot. Mutations are indicated by dark gray shading. **E,** Growth of CO-04-0002 CRC patient-derived xenograft tumors treated with vehicle, KO-2806 (10 mg/kg), MRTX1133 (20 mg/kg), RMC-6236 (25 mg/kg), or KO- 2806-RAS inhibitor doublets for 28 days. Data represent means ± SEM; *n* = 8 mice per group. Statistical significance determined by unpaired Student’s t-test, p-values as indicated on plot. **F,** Growth of GP2D CRC cell-line derived xenograft tumors treated with vehicle, KO-2806 (10 mg/kg), MRTX1133 (10 mg/kg), RMC-6236 (25 mg/kg), or combinations. Each point represents mean tumor volume ± SEM; *n* = 8 mice per group. **G** and **H,** Immunoblots of the indicated signaling proteins in GP2D tumors treated with vehicle, KO-2806, MRTX1133 (**G**), RMC-6236 (**H**) or combinations for 28 days. Doses as indicated in **F**. Images are representative of 2 biological replicates.

Compared to NSCLC cells, mTOR rebound in CRC cells was swifter and more complete, which may be a result of heightened RTK activation in this lineage. Moreover, as this rebound occurs with both mutant-selective and pan-RAS inhibitors, it appears to be independent of wildtype RAS activity, which has been proposed as an adaptive mechanism of escape from mutant-selective KRAS inhibition in NSCLC(13).

To determine whether this improvement in signaling inhibition translates to greater tumor growth inhibition, we treated panels of CRC xenograft models harboring *KRAS^G12D^*or other *KRAS* mutations with MRTX1133 (Fig. 3C) or RMC-6236 (Fig. 3D), respectively. Consistent with our signaling data and results of G12C inhibitor trials, CRC models were generally less sensitive to both mutant-selective and pan-RAS inhibition. Neither MRTX1133 nor RMC-6236 caused regressions in any of the CRC models evaluated when administered as monotherapy. Addition of KO-2806 to either RAS inhibitor improved tumor growth inhibition seen with the single agents, and in some models, induced regressions. *KRAS^G12D^*-mutant models representative of both scenarios were selected for inquiry into the mechanism(s) underpinning their combination responses (Fig. 3E and F, S4C and D). In tumors treated with the combination of MRTX1133 and KO-2806 for 28 days, S6 phosphorylation was potently inhibited compared to tumors treated with MRTX1133 monotherapy (Fig. 3G). Similarly, RMC-6236 monotherapy minimally impacted phospho-S6 levels in tumors (Fig. 3H, S4G), while the combination of RMC-6236 and KO-2806 significantly decreased S6 phosphorylation (Fig. S4E and F). Thus, as was observed with mutant-selective inhibition in *KRAS^G12C^* NSCLC, KO-2806 may deepen antitumor effects of mutant-selective and pan-RAS inhibitors by controlling mTOR activity. Although the combination of KO-2806 and MRTX1133 (Fig. S5A and B) or RMC-6236 (Fig. S5C-H) was well tolerated in mice bearing CRC xenograft tumors, we sought to expand the range of potential dosing schedules ahead of the first-in-human clinical trial. We found that weekly on/off dosing of KO- 2806 resulted in tumor regressions of comparable depth to those observed with continuous dosing (Fig. S6A and B). The same was true of on/off dosing of KO-2806 plus adagrasib in a NSCLC model (Fig. S6C and D), suggesting there may be flexibility in the dose scheduling of these combinations in the clinic.

### Farnesyl transferase inhibition re-sensitizes relapsing tumors to RAS inhibitors

Our data indicate that KO-2806 can improve the sensitivity of treatment-naïve xenograft models to mutant-selective and pan-RAS inhibitors. However, as the patient population treated with KRAS-targeted therapies continues to expand, an ideal combination strategy would benefit patients who have relapsed on a KRAS inhibitor. We therefore asked if KO-2806 may have activity in the context of adaptive resistance to KRAS^G12C^ inhibition. To evaluate this question, we identified a xenograft model derived from a *KRAS^G12C^*-mutant CRC patient that had been treated with adagrasib and a SHP2 inhibitor. We treated this model with adagrasib or the combination of adagrasib and KO-2806 and found tumor growth was better inhibited by the combination than adagrasib monotherapy (Fig. S7A), hinting at the potential for activity in patients with prior exposure to a G12C-selective inhibitor.

We then sought to model NSCLC and CRC tumor progression on RAS inhibitors *in vivo* to encapsulate various potential clinical scenarios. First, we treated *KRAS^G12C^*-mutant NCI-H2122 NSCLC xenograft tumors with adagrasib (Fig. 4A) or sotorasib (Fig. S7B) for 14 days, during which time the tumors steadily increased in volume. After 14 days, half of these animals continued on G12C inhibitor monotherapy, while the other half were administered adagrasib plus KO-2806. Addition of KO-2806 to the dosing regimen resulted in immediate, rapid regression to volumes comparable to those of tumors dosed with the combination from the start of the study. Tumors on G12C inhibitor monotherapy continued to progress, and switching to another inhibitor of the same class (sotorasib to adagrasib) had no effect on progression.

**Figure 4.**
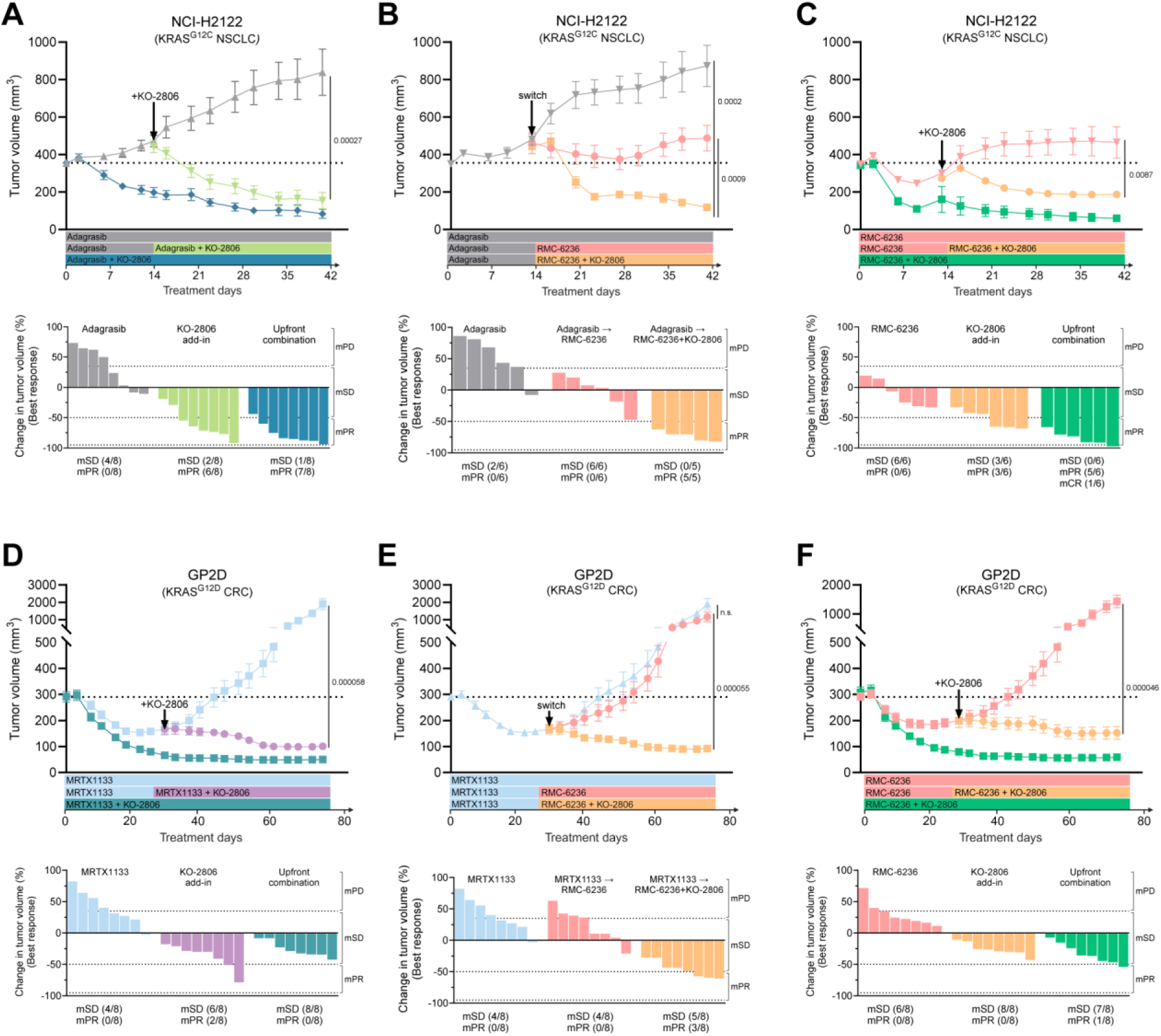
KO-2806 re-sensitizes relapsing tumors to mutant-selective or pan-RAS inhibition. **A**-**C,** Growth of NCI-H2122 xenograft tumors treated with KO-2806 (10 mg/kg PO BID) plus either (**A** and **B**) adagrasib (100 mg/kg PO QD) or (**B** and **C**) RMC-6236 (25 mg/kg PO QD). Single agent and combination treatments were added according to the schedules displayed below tumor volume plots. Data are mean tumor volumes ± SEM, *n =* 5-8 mice per treatment group. Statistical significance between monotherapy group and KO-2806 add-in group was determined by unpaired Student’s t-test, p-values as indicated on plot. Best responses (percent change in tumor volume) and mRECIST response calls were calculated using day 0 as baseline and are displayed below treatment timelines. **D**-**F,** Growth of GP2D xenograft tumors treated with KO-2806 (10 mg/kg PO BID) plus either (**D** and **E**) MRTX1133 (10 mg/kg IP BID) or (**E** and **F**) RMC-6236 (25 mg/kg PO QD) following the schedules displayed below tumor volume plots. Data are mean tumor volumes ± SEM, *n =* 8 mice per treatment group. Statistical significance between groups determined by unpaired Student’s t-test, p-values as indicated on plot. Best responses (percent change in tumor volume) and mRECIST response calls were calculated using the time of KO-2806 add-in (day 28) as baseline and are displayed below treatment timelines.

Signaling analysis revealed that mTOR substrate phosphorylation correlated with antitumor effects (Fig. S7C); mTOR was only well-inhibited in tumors treated with a KO-2806/KRAS^G12C^ inhibitor doublet, implying the mTORC1-centric mechanism of action still applies to the pretreatment setting. We also assessed the impact of KO-2806 add-in on tumors that exhibit an initial response to adagrasib monotherapy. NCI-H2030 tumors regressed on adagrasib alone before regrowing to volumes larger than baseline (Fig. S7D). At that point, KO-2806 addition halted progression, leading to durable tumor stasis.

One candidate therapeutic strategy for patients who progress on KRAS^G12C^(OFF) inhibitors is switching to a RAS inhibitor of another class, as it has been posited patients will not be cross-resistant to (ON) or pan-targeted inhibitors(18). We modeled this scenario in NCI-H2122 tumors by treating them with adagrasib for 14 days before switching to either the pan-RAS(ON) inhibitor RMC-6236, or to the combination of KO-2806 and RMC-6236 (Fig. 4B). Switching inhibitor classes did stop tumor progression and was an improvement compared to adagrasib but switching to the combination of KO-2806 and RMC-6236 elicited significant regressions. For comparison, we also asked if tumors treated with RMC-6236 from the start could still benefit from addition of KO-2806. RMC-6236 administered up-front was superior to KRAS^G12C^(OFF) inhibitors, resulting in tumor stasis, but KO-2806 addition after 14 days led to rapid tumor regression (Fig. 4C). Together these findings imply that KO-2806 can resensitize *KRAS^G12C^*- mutant NSCLC tumors to RAS inhibition following progression on a RAS inhibitor, regardless of inhibitor class.

To ask if KO-2806 similarly overcomes adaptive resistance in CRC, we modeled the equivalent scenarios in a *KRAS^G12D^*-mutant CRC xenograft model. The G12D-selective inhibitor MRTX1133 induced GP2D tumor regression for approximately 3 weeks before relapse (Fig. 4D). Adding KO-2806 to the treatment regimen of relapsing tumors at 4 weeks led to deeper tumor regression, comparable to up-front administration of KO-2806 and MRTX1133. We next asked if switching RAS inhibitor class during relapse would control growth of this CRC model compared to a KO-2806/RAS inhibitor combination. Switching to single agent RMC-6236 treatment had no effect on the growth of tumors relapsing on MRTX1133 (Fig. 4E). This result stands in contrast to our findings in the NSCLC model, where RMC-6236 induced tumor stasis after progression on adagrasib, suggesting that wildtype RAS may play less of a role in adaptive resistance to KRAS inhibition in CRC compared to NSCLC(12, 19). Switching GP2D tumors to the combination of KO-2806 and RMC-6236 reversed relapse and resulted in further tumor regression. Finally, we asked whether KO-2806 addition could rescue tumors progressing on RMC-6236 administered up-front. Again, in contrast to NSCLC, RMC-6236 was not superior to mutant-selective MRTX1133 and GP2D tumors regressed and then relapsed on a similar timeline. Adding KO-2806 to RMC-6236 upon relapse once again led to tumor regression, similar to up-front administration of this combination.

Combinations of KO-2806 and adagrasib, sotorasib, RMC-6236, or MRTX1133 were well tolerated in lengthy studies ranging from 42 to 77 days (Fig. S8). As such, these results indicate that addition of KO-2806 tolerably and effectively reverses relapse of NSCLC and CRC xenograft models progressing on multiple classes of RAS inhibitors.

## DISCUSSION

Although inactive-state KRAS^G12C^ inhibitors have generated meaningful results in the clinic, resistance remains a major problem limiting their therapeutic benefit. Mechanisms underlying clinical resistance are heterogeneous but ultimately converge on reactivating MAPK pathway or compensatory PI3K-AKT-mTOR signaling(6, 7). Because these inhibitors only target inactive, GDP-bound KRAS, tumor cells retain a reasonable window to restore oncogenic signaling through adaptive or acquired means. In this study, we show that mTORC1 plays a critical role in limiting response to KRAS^G12C^(OFF) inhibitors, providing mechanistic rationale for combining adagrasib with the farnesyl transferase inhibitor KO-2806, a combination being evaluated clinically as part of the first-in-human FIT-001 trial (NCT06026410).

Genomic analyses of *KRAS^G12C^*-mutant cancers that progressed on adagrasib after exhibiting initial response or disease stabilization point to genetic reactivation of MAPK signaling as the predominant driver of acquired resistance to inactive-state G12C inhibition(9, 20). Moreover, preclinical studies have outlined a mechanism of adaptive resistance to mutant-selective KRAS inhibitors whereby EGFR drives GTP-loading of mutant KRAS and/or activation of wildtype RAS isoforms, restoring MAPK signaling(12, 13). These findings provide rationale for the use of active-state inhibitors (KRAS-mutant selective or pan-RAS) and support combination of KRAS inhibitors with anti-EGFR antibodies, SHP2 inhibitors, and SOS1 inhibitors(21, 22). Still, the one-third of patients who progressed on adagrasib without an identifiable acquired genetic driver of resistance, and the documented impact of co-mutations such as *STK11* and *PIK3CA* on therapy response, hint at the existence of MAPK-independent escape mechanisms. Based on the consistency in which we identified persistent mTORC1 activity in models partially or minimally sensitive to RAS inhibition, it is unlikely that therapeutic strategies designed to block MAPK reactivation will suffice in all cases. Our results imply that failure to control mTOR in tumors with any degree of dependence on this pathway (due to cell lineage, levels of RTK activity, co-mutations) leaves an opportunity for rapid escape from RAS-targeted therapies. KO- 2806 blocks this escape route by inhibiting RHEB-dependent mTORC1 activity, making it an attractive potential combination partner to overcome resistance to RAS inhibitors.

We began this study looking to demonstrate enhanced benefit from up-front administration of KO-2806 and a KRAS inhibitor over the KRAS inhibitor alone in xenograft models.

However,rapid development of diverse approaches to inhibit mutant KRAS calls for therapeutic interventions that may benefit patients who have had prior exposure to RAS inhibitors. We modeled progression of NSCLC and CRC tumors on mutant-selective and pan-RAS inhibitors and found that KO-2806 addition during progression caused rapid and durable tumor regression in all assessed xenograft models. These results are consistent with the notion that mTORC1 represents an “easy” escape route for *KRAS*-mutant tumors, and that overcoming simultaneous MAPK and mTOR blockade is more challenging. It has been demonstrated that active-state RAS inhibitors have preclinical and clinical activity in models/patients that failed on a KRAS^G12C^(OFF) inhibitor(23, 24). In our NSCLC preclinical model, we indeed found switching to RMC-6236 slowed tumor progression compared to remaining on adagrasib. Nonetheless, adding KO-2806 to adagrasib was superior to switching to single agent RMC-6236. In contrast, in our CRC preclinical model, switching to RMC-6236 was indistinguishable from remaining on mutant-selective MRTX1133. This result suggests that reactivation of mutant and/or wildtype RAS may play a larger role in mediating resistance in NSCLC compared to in CRC. Nevertheless, only addition of KO-2806 induced tumor regression following progression on a RAS inhibitor, establishing mTORC1 as a valuable player in RAS inhibitor tolerance. Many prior inquiries cite ERK reactivation as a major impediment to KRAS inhibitor efficacy and propose vertical inhibition of the MAPK pathway as a strategy to overcome resistance. This work identifies an alternative route of therapy escape via mTORC1 and nominates a combination approach to address this. In our view, tackling adaptive and acquired resistance to RAS inhibitors will likely require informed application of both strategies. The challenge lies in defining the appropriate context(s) in which to apply each approach. Future preclinical studies in models derived from patients with prior exposure to RAS inhibitors, and isogenic models generated based on known genetic drivers of resistance may help to do so. Clinical biomarker analyses, including retrospective efforts and in-trial monitoring via liquid biopsies, may be considered to anticipate a particular tumor’s route of therapy evasion and to design therapeutic intervention accordingly.

As additional RAS-targeted agents advance to approval, the population of *KRAS*-mutant patients that have seen one or more RAS inhibitors will expand. Here, we have demonstrated that the farnesyl transferase inhibitor KO-2806 can expand the utility of multiple classes of RAS inhibitors in tumors that develop or intrinsically possess resistance to these agents. By targeting mTORC1 through inhibition of RHEB farnesylation, KO-2806 shuts down a vital survival pathway for many tumors, re-sensitizing them to RAS inhibition. Therefore, the combination of KO-2806 and RAS inhibitors represents a versatile and promising therapeutic strategy for patients, irrespective of prior RAS inhibitor therapy received.

## METHODS

### Synthesis of KO-2806

KO-2806 may be synthesized using methods described in PCT Patent Application Publication No. WO2024220600A1.

#### Reagents and Cell Lines

All compounds were either synthesized at WuXi AppTec or purchased from MedChemExpress. Compounds were formulated in 100% DMSO for *in vitro* studies. T24, NCI-H2122, NCI-H2030, LoVo, and HCT116 cell lines were obtained from American Type Culture Collection (ATCC) and the GP2D cell line was obtained from Creative Bioarray. T24 and HCT116 cells were maintained in McCoy 5A + 10% heat-inactivated fetal bovine serum (FBS) supplemented with 1.5 mM L-glutamine, 25 mM HEPES, 100 U/mL penicillin, and 100 U/mL streptomycin (Gibco). NCI-H2122 and NCI-H2030 cells were maintained in RPMI-1640 + 10% heat-inactivated FBS supplemented with 4.5 g/L D-glucose, 2.383 g/L HEPES, 2 mM L-glutamine, 1.5 g/L sodium bicarbonate, 110 mg/L sodium pyruvate, 100 U/mL penicillin, and 100 U/mL streptomycin (Gibco). GP2D cells were maintained in DMEM + 10% heat-inactivated FBS supplemented with 4.5 g/L D-glucose, 4 mM L-glutamine, 100 U/mL penicillin, and 100 U/mL streptomycin (Gibco). LoVo cells were maintained in F-12K + 10% heat-inactivated FBS supplemented with 2 mM L-glutamine, 100 U/mL penicillin, and 100 U/mL streptomycin (Gibco). Cell lines were maintained in a humidified incubator at 37°C, 5% CO_2_ and tested negative for mycoplasma.

#### Biochemical Activity Assays

FTase and GGTase activity were measured using fluorescence polarization assay (WuXi AppTec, Shanghai). Based on the assay, recombinant human FTase (2.5 nM) or GGTase (25 nM), dansyl labeled GCVLS peptide (85 nM) or GCVLL peptide (10 μM), FFP (30 nM) or GGPP (10 μM), and KO-2806 (10 mM stock in DMSO) were mixed in assay buffer (50 mM Tris, pH 7.5, 10 mM magnesium chloride, 10 μM zinc chloride, 0.08% CHAPS, 5 mM dithiothreitol) and incubated in a total volume of 20 μL at 30°C for 50-120 minutes. Fluorescence intensity was detected (excitation at 340 nm and emission at 486 nM) using EnVision (PerkinElmer; Waltham, MA).

#### Cell Proliferation Assay

The ability of KO-2806 to inhibit cell proliferation was evaluated (WuXi AppTec, Shanghai). T24/83 cells were seeded at 100 cells/well in 384 well plates and grown in the presence of the indicated concentrations of KO-2806 (20 mM stock in DMSO), 0.7% DMSO as a negative control, or 100 μM tipifarnib as a positive control at 37°C for 4 days. Cell viability was assessed using the CellTiter-Glo kit (Promega; Madison, WI). Luminescence was recorded using an EnVision microplate reader (PerkinElmer; Waltham, MA).

#### Spheroid Growth Assay

Cells were seeded in 96-well ultralow attachment plates at a density of 2,000 cells/well. Cells were centrifuged at 1000 rpm for 2 min to form spheroids. The following day, spheroids were treated with DMSO, KO-2806, adagrasib, or the combination. Spheroids were incubated at 37°C for 0 (baseline), 3, and 5 days. Spheroid growth was assessed using CellTiter-Glo 3D kit (Promega; Madison, WI). Luminescence was recorded using Spark microplate reader (Tecan; Switzerland).

#### Transfection of siRNA

NCI-H2122 cells (500,000) were seeded on 60 mm dishes and transfected with Dharmacon ON-TARGETplus siRNA SMARTPool against RHEB (Horizon Discovery) or a non-targeting pool, used as a negative control, using Lipofectamine RNAiMAX (ThermoFisher). Cells were incubated for 72 hours prior to collection for immunoblotting to allow for depletion of *RHEB* expression. These siRNA-transfected NCI-H2122 cells were used for downstream applications.

#### Stable Cell Line Generation

NCI-H2122 cells were stably transduced with lentivirus vectors encoding HA-tagged RHEB wildtype (WT) or C181S farnesylation-deficient mutant (Transomic). Polybrene was supplemented to improve transduction. After 24 hours, the media was replaced with fresh growth medium. After another 24 hours, transduced cells were selected with 10 µg/mL blasticidin for 7 days. These stable cell lines were then transduced with a Tet-On 3G lentivirus vector encoding a doxycycline inducible shRNA with the 3’ UTR targeting sequence against endogenous RHEB (Transomic). Cells were selected with 2 µg/mL puromycin in the appropriate medium supplemented with tetracycline-free FBS for 5 days. These cells were maintained in selection medium and treated with 1 µg/mL doxycycline for downstream applications.

#### Immunoblotting

For immunoblotting of cell lines grown in 2D, 2 million cells were plated onto 10 cm dishes. For 3D immunoblotting, 10,000 cells/well were plated onto 96-well ultralow attachment plates and centrifuged at 1000 rpm for 2 min to form spheroids. 96 spheroids were pooled together for one sample. Cell lysates were prepared on ice by washing cells once with PBS, resuspending in 1X cell lysis buffer (Cell Signaling Technology #9803) or RIPA buffer supplemented with Halt protease inhibitor cocktail (Thermo Scientific #78430) and briefly sonicating or vortexing. Tumor lysates were prepared by adding tumor fragments to hard tissue homogenizing tubes (2 mL reinforced polypropylene tubes) containing RIPA buffer and five 2.8 mm ceramic beads (Fisher Scientific). Tumors were then homogenized for 30s at 4.5 m/s using a bead mill. Lysates were cleared by centrifugation (maximum speed, 10 min) and protein concentration was determined by BCA assay (Pierce). 20-60 µg of lysate was loaded on to 4-12% Bis-Tris gels (NuPAGE, Invitrogen) for electrophoresis and immunoblotting.

#### Immunofluorescence (IF) Imaging

NCI-H2122 cells with stably expressed HA-tagged RHEB WT or C181S-mutant were seeded in 8-well chamber slides (ibidi) at a density of 5,000 cells/well. Cells were treated with either DMSO or 1 μM KO-2806 for 24 hours. Cells were fixed with 4% PFA in PBS for 10 minutes at room temperature, then washed 3 times with PBS. Cells were permeabilized in 0.1% Triton X- 100 in PBS for 5 minutes, washed 3 times with PBS, and incubated in blocking buffer (1% BSA + 0.01% Tween 20 in PBS) for 1 hour at room temperature. Cells were labeled with HA-tag rabbit mAb Alexa Fluor 488 conjugate (Cell Signaling Technology) at 1:50 dilution in blocking buffer for 1 hour at room temperature in the dark. Following 3 washes with PBS, mounting medium with DAPI (ibidi) was added on the cells for 10 minutes at room temperature in the dark. IF images were taken on a Keyence microscope with 60X oil objective.

### In Vivo Studies

All xenograft studies were conducted at Crown Bioscience or WuXi AppTec. All procedures involving the care and use of animals in these studies were reviewed and approved by the Institutional Animal Care and Use Committee (IACUC) at each facility prior to execution. During the study, the care and use of animals were conducted in accordance with the regulations of the Association for Assessment and Accreditation of Laboratory Animal Care (AAALAC). Female, 4- 8 weeks old, BALB/c nude or NOD/SCID mice were used. The following human cancer cell line-derived xenograft (CDX) models were used for these studies: NCI-H2122, NCI-H2030, NCI- H358, NCI-H1792, LU99, NCI-H23, SW1573, GP2D, SW480, HCT116, SW620, LoVo, and DLD-1. The following human cancer patient-derived xenograft (PDX) models were used for these studies: HN1420, LU2512, LU11693, LU6405, CO-04-0002, CR1245, CR3262, CR3150, and CR9528.

To generate CDX models, tumors cells were maintained *in vitro* with appropriate growth medium supplemented with 10% fetal bovine serum at 37 °C in an atmosphere of 5% CO_2_ in air. The cells in exponential growth phase were harvested and quantitated by cell counter before tumor inoculation. Unless otherwise specified, each mouse was inoculated subcutaneously in the right upper flank region with 0.5-2×10^7^ in 0.1-0.2 mL of PBS tumor cells per mouse mixed with Matrigel (1:1) for tumor development. To generate PDX models, tumor fragments from stock mice were harvested and used for inoculation into mice. Each mouse was inoculated subcutaneously in the right upper flank with primary human tumor xenograft model tumor fragment (2-3 mm in diameter) for tumor development.

The randomization started when the mean tumor size reached approximately 200-400 mm^3^. Randomization was performed based on “Matched distribution” method (StudyDirector^TM^ software, version 3.1.399.19). The date of randomization was denoted as Day 0. The treatment was initiated on the same day of randomization (Day 0) as per the study design. After tumor inoculation, the animals were checked daily for morbidity and mortality. During routine monitoring, the animals were checked for any effects of tumor growth and treatments on behavior such as mobility, food and water consumption, body weight gain/loss (body weights would be measured three times/daily per week after randomization), eye/hair matting and any other abnormalities. Mortality and observed clinical signs were recorded for individual animals in detail. Tumor volumes were measured three times per week after randomization in two dimensions using a caliper, and the volume was expressed in mm^3^ using the formula: V = (L x W x W)/2, where V is tumor volume, L is tumor length (the longest tumor dimension), and W is tumor width (the longest tumor dimension perpendicular to L). Dosing as well as tumor and body weight measurements were conducted in a laminar flow cabinet. The body weights and tumor volumes were measured by using Study Director ^TM^ software (version 3.1.399.19).

Xenograft mice were treated by oral gavage with vehicle, KO-2806 in aqueous formulation at indicated doses twice a day, adagrasib (in 10% research grade Captisol in 50 mM citrate buffer, pH 5.0) at indicated doses daily, sotorasib (in 2% HPMC and 1% Tween 80) at indicated doses daily, or RMC-6236 (in 10% research grade Captisol in 50 mM citrate buffer, pH 5.0) as well as the various combinations. Xenograft mice were treated by intraperitoneal administration with MRTX1133 (in 10% Captisol, 50 mM citrate buffer, pH 5.0) at indicated doses twice a day as well as the combination with KO-2806. The treatment period was performed for 4-8 weeks. For studies with tumor sample collections, each tumor was split into 2 pieces; half was snap frozen for protein and the other half fixed for histological studies.

Statistical analysis of the difference in mean tumor volume between the single agent KRAS-mutant selective or pan-RAS inhibitor-treated group and the KO-2806 combination-treated group was calculated using a two-tailed unpaired Student’s t-test in GraphPad Prism with p-value < 0.05 considered statistically significant. The percentage tumor volume change from specified baseline for individual xenograft mice was graphed in waterfall plots, and best response calls were made using mRECIST criteria(25).

#### Immunohistochemistry (IHC)

All immunostaining was performed at Histowiz, Inc., Brooklyn, New York, using the Leica Bond RX automated stainer (Leica Microsystems). The slides were dewaxed using xylene and alcohol based dewaxing solutions. Epitope retrieval was performed by heat-induced epitope retrieval (HIER) of the formalin-fixed, paraffin-embedded (FFPE) tissue using citrate-based pH 6 solution for 20 mins at 95 °C. The tissues were first incubated with peroxide block buffer (Leica Microsystems), followed by incubation with the primary antibodies Ki67 (ab15580), Cleaved Caspase 3 (CST9661), p-S6 (CST4858), or p-4EBP1 (CST2855) at 1:800, 1:300, 1:200, and 1:800 dilutions respectively for 30 mins, followed by DAB rabbit secondary reagents: polymer, DAB refine and hematoxylin (Leica Microsystems). The slides were dried, cover slipped and visualized using a Leica Aperio AT2 slide scanner (Leica Microsystems). HALO was used for quantitative image analysis.

## Data Availability

Data are available from the corresponding author upon reasonable request.

## Supplementary data

**Figure S1.**
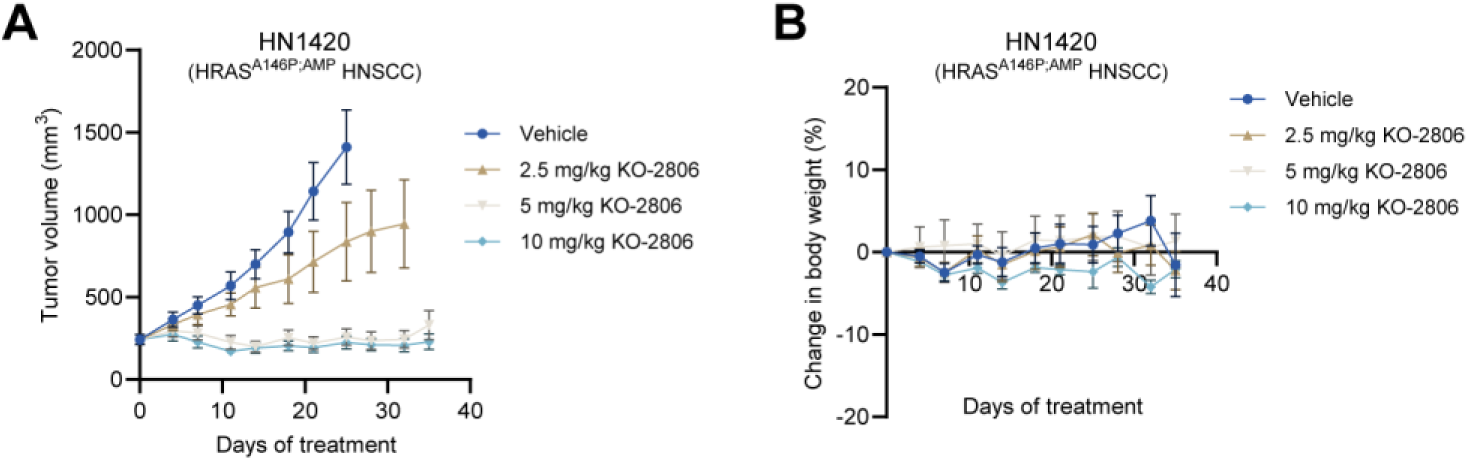
KO-2806 inhibits growth of HRAS-mutant head and neck patient-derived xenograft tumors. **A**, Growth of HN1420 PDX tumors treated with indicated doses of KO-2806 PO BID for 35 days. Data are mean tumor volumes ± SEM, *n =* 6 mice per group. **B**, Percent change in body weight of animals from (**A**). Data are means ± SEM, *n =* 6 mice per group.

**Figure S2.**
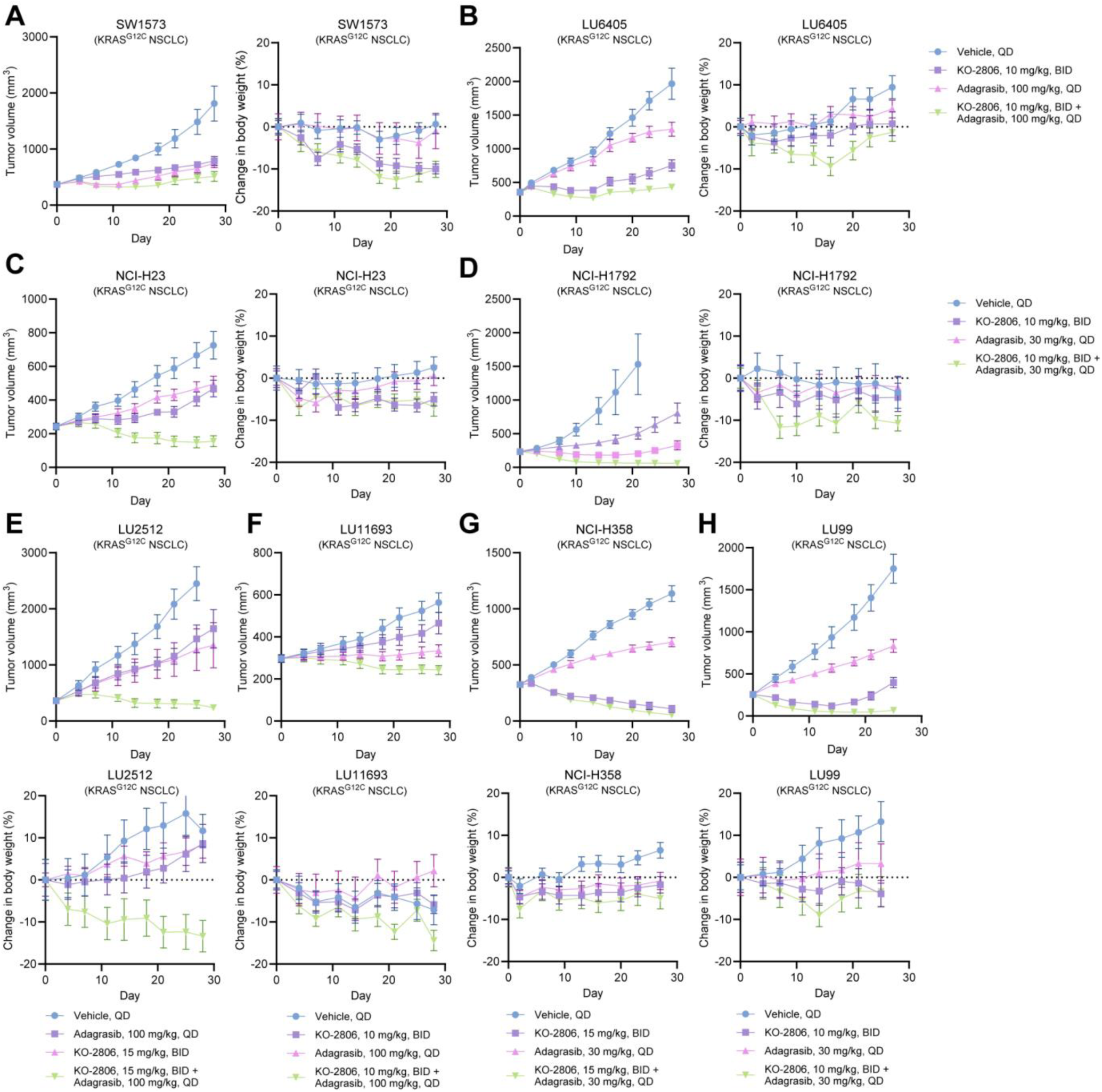
Antitumor efficacy and tolerability of combined adagrasib and KO-2806 in *KRAS^G12C^*-mutant NSCLC xenograft models. **A**-**H**, Tumor volumes and percent change in body weight of xenograft-bearing mice treated with the indicated doses of KO-2806 and/or adagrasib by oral gavage for 28-35 days. Data are means ± SEM, *n* = 6-8 mice per treatment group.

**Figure S3.**
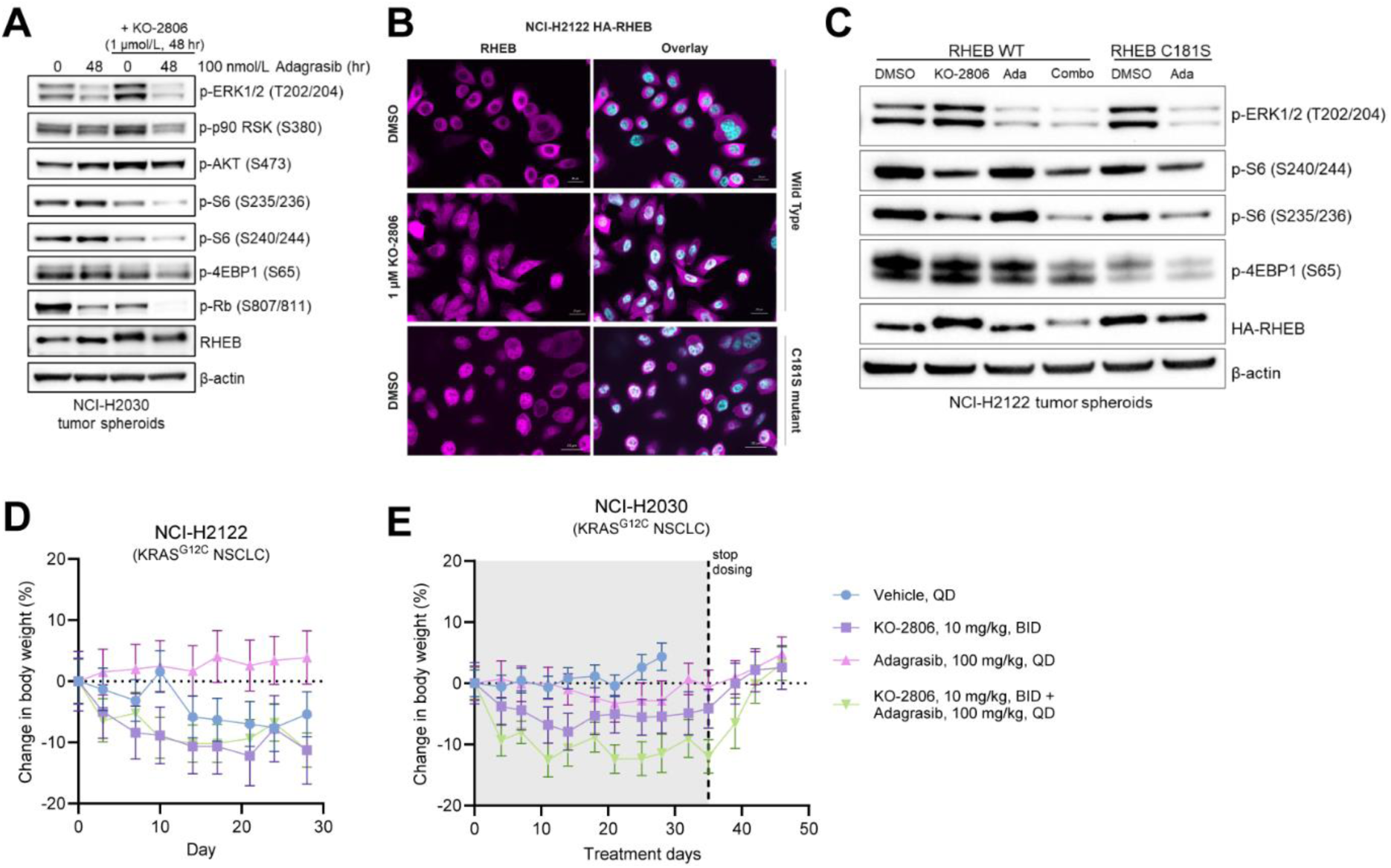
Combination of KO-2806 and adagrasib inhibits mTOR signaling and is tolerable *in vivo*. **A**, Immunoblots of the indicated signaling proteins in NCI-H2030 spheroids exposed to 100 nmol/L adagrasib for 0 or 48 hours in the presence or absence of 1 μmol/L KO-2806. Images are representative of 2 biological replicates. **B**, Immunofluorescence images of NCI-H2122 cells expressing HA-tagged RHEB wildtype or C181S unfarnesylated mutant in the presence or absence of 1 μmol/L KO-2806 for 24 hours. Localization of RHEB (magenta) was visualized using nuclear DAPI stain (blue). Images are representative of 2 biological replicates. Scale bar, 50 μm. **C**, Immunoblots of the indicated signaling proteins in NCI-H2122 spheroids expressing HA-tagged RHEB wildtype or C181S mutant exposed to 100 nmol/L adagrasib in the presence or absence of 1 μmol/L KO-2806 for 48 hours. Images are representative of 2 biological replicates. **D** and **E**, Percent change in body weight of mice bearing NCI- H2122 (**D**) or NCI-H2030 (**E**) xenograft tumors treated with KO-2806, adagrasib, or the combination for 28-35 days. Data are means ± SEM, *n* = 8 mice per treatment group.

**Figure S4.**
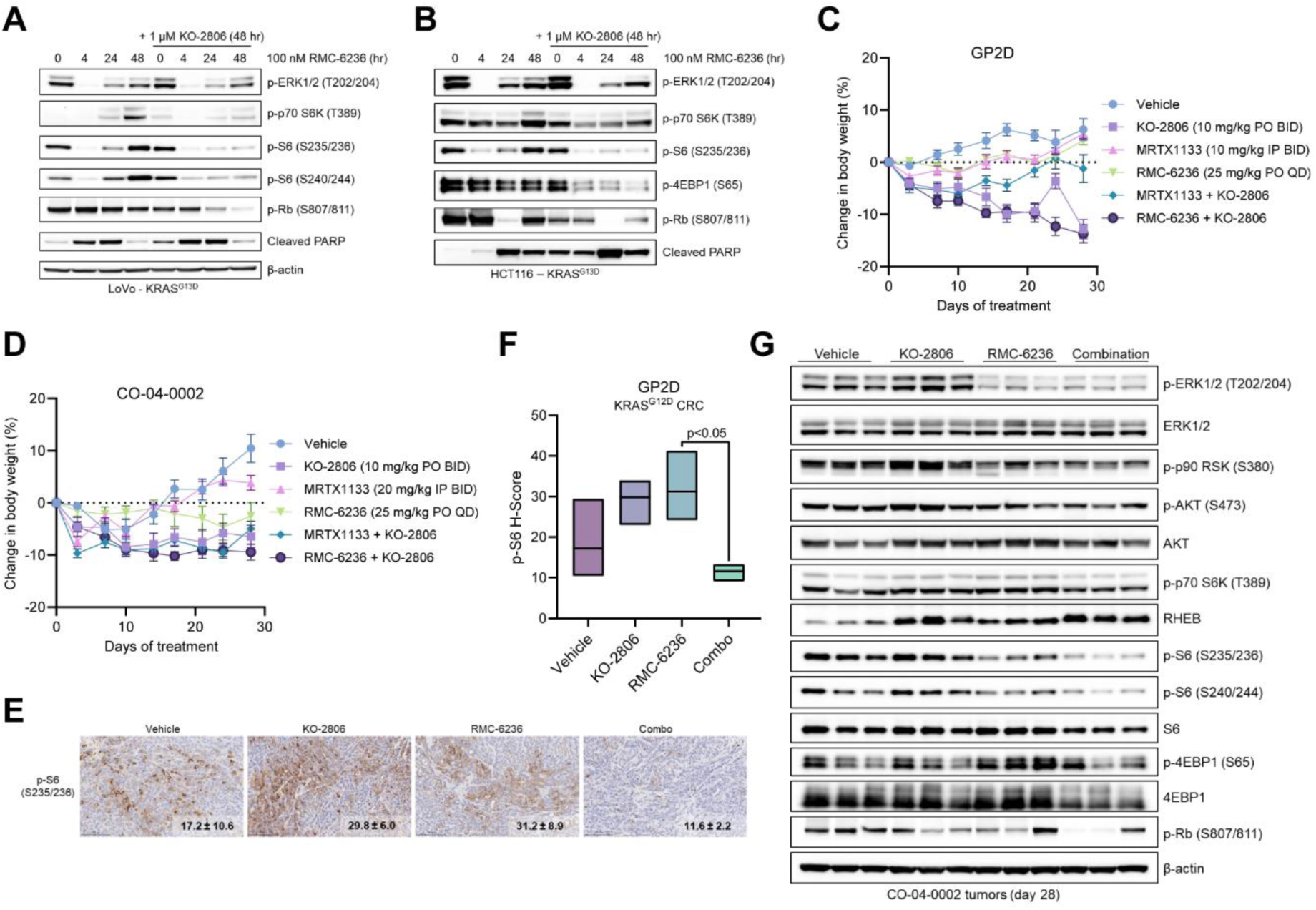
The combination of KO-2806 and mutant-selective KRAS or pan-RAS inhibitors inhibits mTOR signaling in CRC models and is tolerable *in vivo*. **A** and **B**, Immunoblots of the indicated signaling proteins in LoVo (**A**) or HCT116 (**B**) G13D-mutant colorectal cancer cells exposed to 100 nmol/L RMC-6236 for 0, 4, 24, or 48 hours in the presence or absence of 1 μmol/L KO-2806. Images are representative of 2 biological replicates. **C** and **D**, Percent change in body weight of mice bearing GP2D (**C**) or CO-04-0002 (**D**) xenograft tumors treated with the indicated doses of KO-2806, MRTX1133, RMC-6236, or combinations. Data are mean percent changes in body weight ± SEM, *n* = 8 mice per group. **E** and **F**, Immunohistochemistry analysis of GP2D tumors treated with vehicle, KO-2806 (10 mg/kg PO BID), RMC- 6236 (25 mg/kg PO QD), or the combination for 28 days. Images (**E**) are representative of 3 tumors. Scale bar, 100 μm. Quantification of phospho-S6 staining (**F**) presented as mean H-scores ± SD, *n* = 3 tumors. **G**, Immunoblots of the indicated signaling proteins in CO-04-0002 CRC patient-derived xenograft tumors treated with vehicle, KO-2806 (10 mg/kg), RMC-6236 (25 mg/kg), or the combination for 28 days.

**Figure S5.**
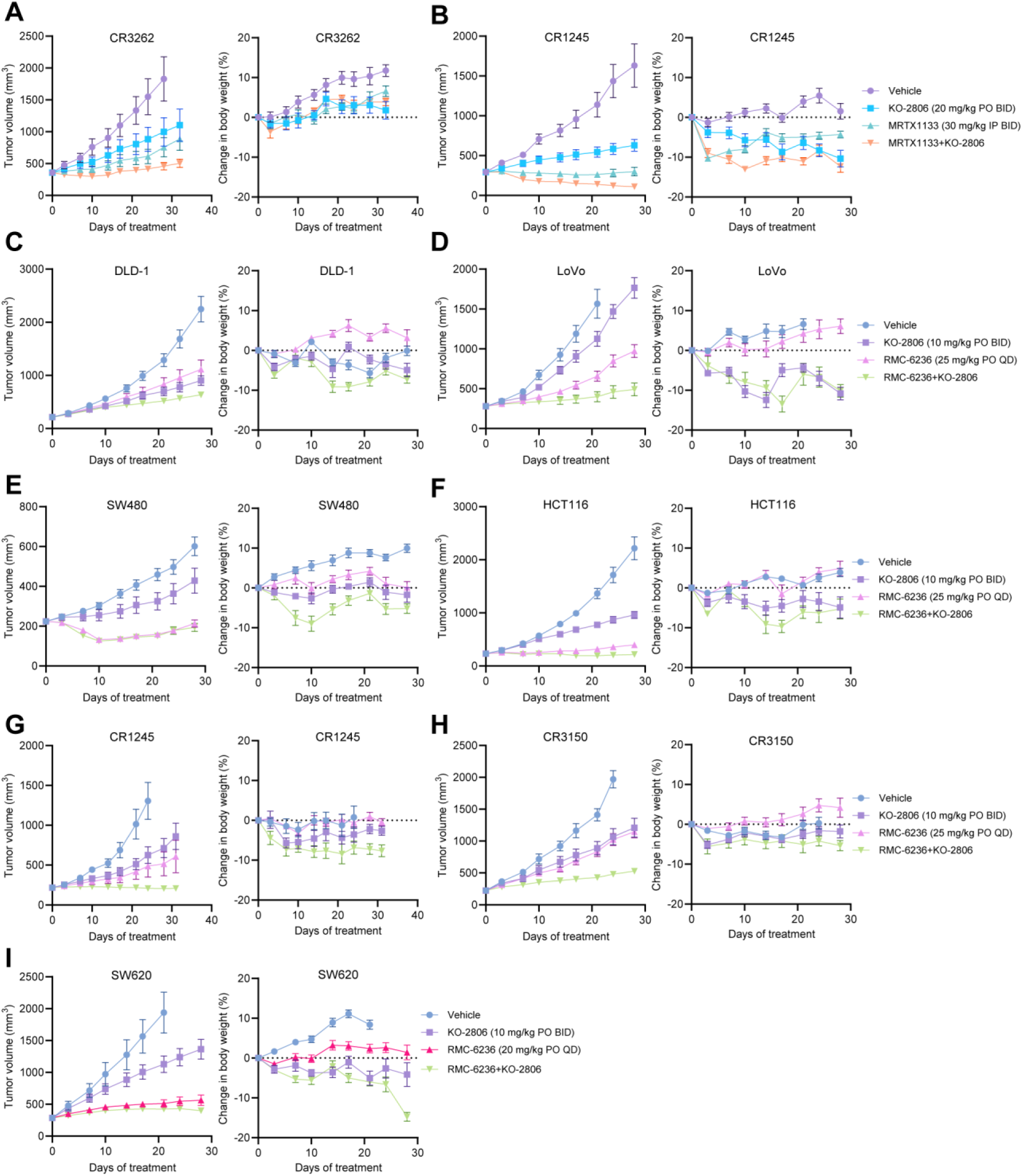
Antitumor efficacy and tolerability of combined KO-2806 and MRTX1133 or RMC-6236 in *KRAS*-mutant CRC xenograft models. **A**-**I**, Tumor volumes and percent change in bodyweight of xenograft-bearing mice treated with the indicated doses of KO-2806, MRTX1133, or RMC-6236 for 28-35 days. Data are means ± SEM, *n* = 8 mice per treatment group.

**Figure S6.**
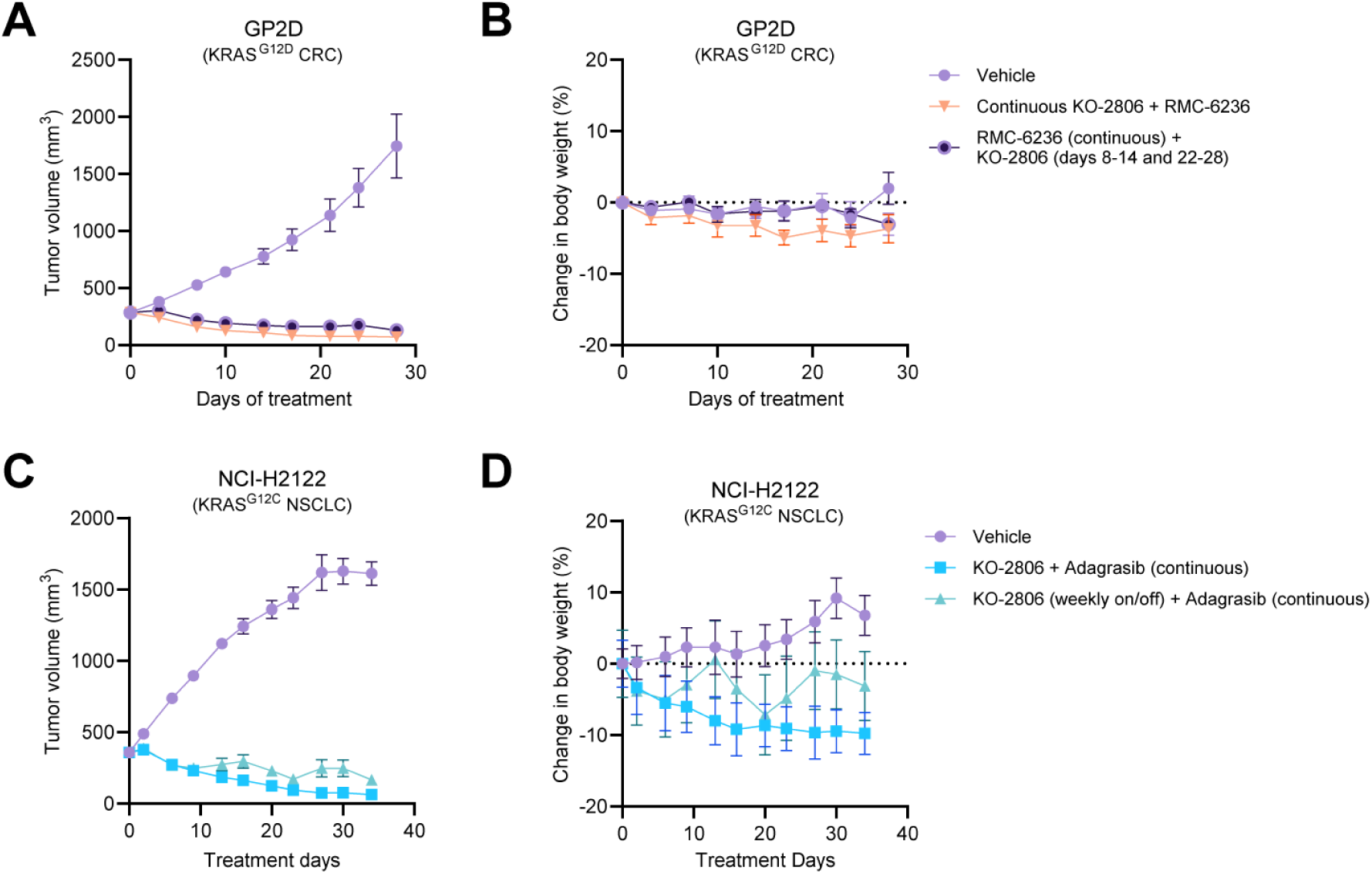
Intermittent dosing of KO-2806 leads to tumor regressions in combination with RAS inhibitors. **A** and **B**, Tumor volume (**A**) and percent change in body weight (**B**) of mice bearing GP2D xenograft tumors treated with vehicle or 25 mg/kg RMC-6236 PO QD plus 10 mg/kg KO-2806 PO BID either continuously (days 0-28) or on/off weekly (treatment on days 8-14 and 22-28). **C** and **D**, Tumor volume (**C**) and percent change in body weight (**D**) of mice bearing NCI-H2122 xenograft tumors treated with vehicle or 100 mg/kg adagrasib PO QD plus 10 mg/kg KO-2806 PO BID either continuously (days 0-35) or on/off weekly (treatment on days 0-7, 15-21, 29-35).

**Figure S7.**
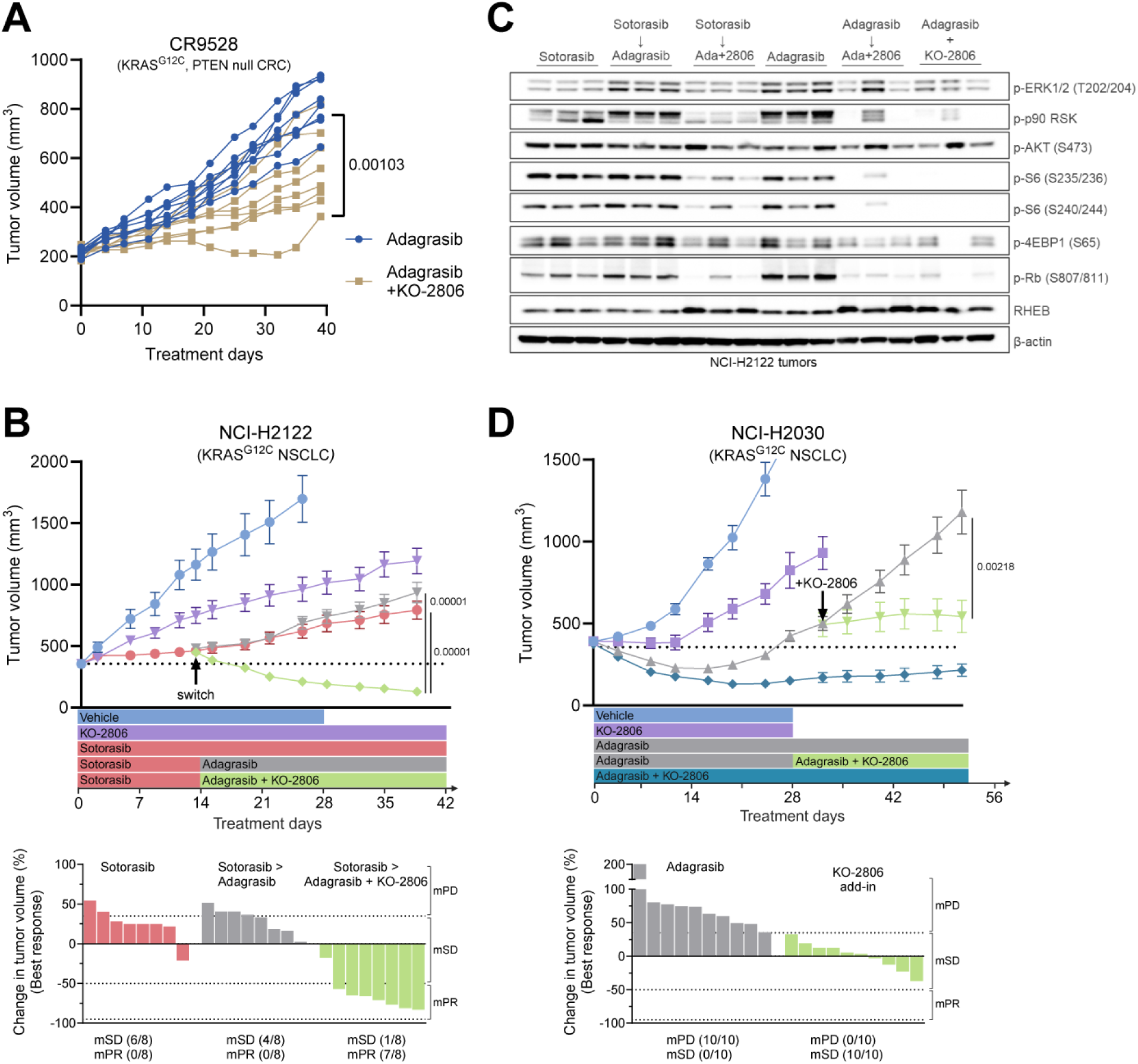
KO-2806 addition re-sensitizes NSCLC tumors to KRAS^G12C^ inhibition. **A**, Growth of individual CR9528 xenograft tumors derived from a CRC patient who had been treated with adagrasib and a SHP2 inhibitor. Xenograft tumors were treated with 100 mg/kg adagrasib PO QD or adagrasib plus 10 mg/kg KO-2806 PO BID for 41 days, *n =* 8 mice per group. Statistical significance determined by unpaired Student’s t-test, p-value as indicated on plot. **B**, NCI-H2122 xenograft tumors were treated with 10 mg/kg KO-2806, 100 mg/kg sotorasib, 100 mg/kg adagrasib, or the combination of KO-2806 plus adagrasib for 41 days total. Treatments were added according to the schedules displayed below tumor volume plots. Data are means ± SEM, *n =* 5-8 mice per group. Statistical significance determined by unpaired Student’s t-test, p-values as indicated on plot. Best responses (percent change in tumor volume) and mRECIST response calls were calculated using day 0 as baseline and are displayed below treatment timelines. **C**, Immunoblots of the indicated signaling proteins in NCI-H2122 tumors treated with sotorasib, adagrasib, KO-2806, or the combinations according to the timelines in Fig. 4A and S7B. **D,** NCI-H2030 xenograft tumors were treated with 10 mg/kg KO-2806, 100 mg/kg adagrasib, or the combination for 52 days. Treatments were added according to the schedules displayed below tumor volume plots. Data are means ± SEM, *n =* 5-10 mice per group. Statistical significance determined by unpaired Student’s t-test, p-values as indicated on plot. Best responses (percent change in tumor volume) and mRECIST response calls were calculated using day 28 as baseline and are displayed below treatment timelines.

**Figure S8.**
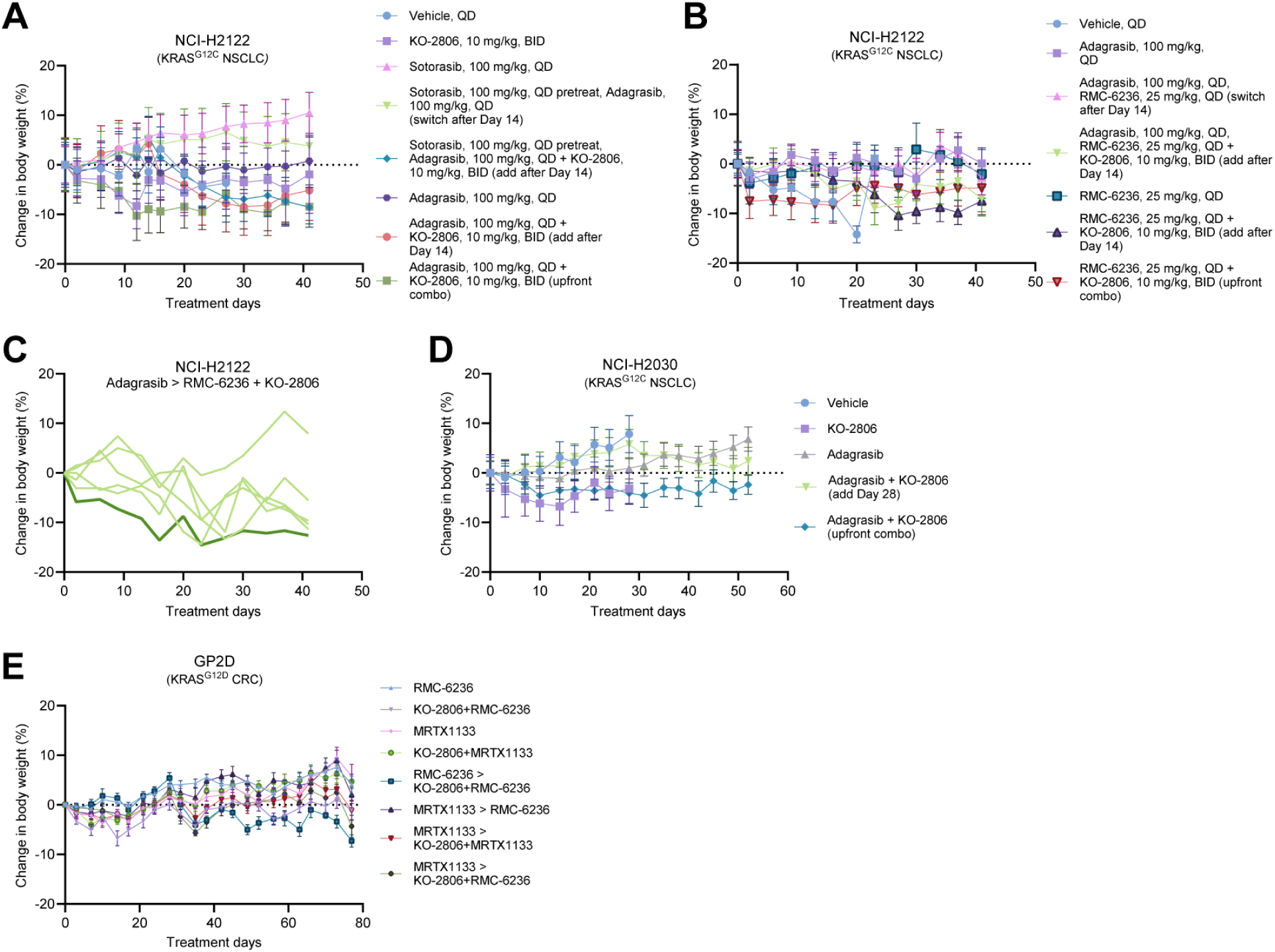
Prolonged treatment with KO-2806/RAS inhibitor combinations is well tolerated in mice. **A** and **B**, Percent change in body weight of mice bearing NCI-H2122 xenograft tumors treated with KO-2806, adagrasib, (**A**) sotorasib, (**B**) RMC-6236, or the combinations as specified for 41 days. Data are mean percent change in body weight ± SEM, *n =* 6-8 mice per group. **C**, Percent change in body weight of individual mice from indicated group in **B** with emphasis on one animal (dark line) excluded from tumor volume mean in Fig. 4B due to extended dosing holidays. **D**, Percent change in body weight of mice bearing NCI-H2030 xenograft tumors treated with KO-2806, adagrasib, or the combination as specified for 52 days. Data are mean percent change in body weight ± SEM, *n =* 5-10 mice per treatment group. **E**, Percent change in body weight of animals treated with 25 mg/kg RMC-6236 PO QD, 10 mg/kg KO-2806 PO BID, 10 mg/kg MRTX1133 IP BID, or the combinations. Data are mean percent change in body weight ± SEM, *n =* 8 mice per group.

## Notes

### Competing Interest Statement

HVP, AES, SC, JGG, LK, AM, YAL, FB, and SM are employees and stockholders of Kura Oncology. TS and XZ are former employees of Kura Oncology.

## REFERENCES

1. Hallin J, Engstrom LD, Hargis L, Calinisan A, Aranda R, Briere DM, et al. The KRAS(G12C) Inhibitor MRTX849 Provides Insight toward Therapeutic Susceptibility of KRAS- Mutant Cancers in Mouse Models and Patients. Cancer Discov. 2020;10(1):54–71.

2. Canon J, Rex K, Saiki AY, Mohr C, Cooke K, Bagal D, et al. The clinical KRAS(G12C) inhibitor AMG 510 drives anti-tumour immunity. Nature. 2019;575(7781):217-23.

3. Hallin J, Bowcut V, Calinisan A, Briere DM, Hargis L, Engstrom LD, et al. Anti-tumor efficacy of a potent and selective non-covalent KRAS(G12D) inhibitor. Nat Med. 2022;28(10):2171–82.

4. Jiang J, Jiang L, Maldonato BJ, Wang Y, Holderfield M, Aronchik I, et al. Translational and Therapeutic Evaluation of RAS-GTP Inhibition by RMC-6236 in RAS-Driven Cancers. Cancer Discov. 2024;14(6):994–1017.

5. Singhal A, Li BT, O’Reilly EM. Targeting KRAS in cancer. Nat Med. 2024;30(4):969–83.

6. Lietman CD, Johnson ML, McCormick F, Lindsay CR. More to the RAS Story: KRAS(G12C) Inhibition, Resistance Mechanisms, and Moving Beyond KRAS(G12C). Am Soc Clin Oncol Educ Book. 2022;42:1-13.

7. Zhu C, Guan X, Zhang X, Luan X, Song Z, Cheng X, et al. Targeting KRAS mutant cancers: from druggable therapy to drug resistance. Mol Cancer. 2022;21(1):159.

8. Kitai H, Choi PH, Yang YC, Boyer JA, Whaley A, Pancholi P, et al. Combined inhibition of KRAS(G12C) and mTORC1 kinase is synergistic in non-small cell lung cancer. Nat Commun. 2024;15(1):6076.

9. Zhao Y, Murciano-Goroff YR, Xue JY, Ang A, Lucas J, Mai TT, et al. Diverse alterations associated with resistance to KRAS(G12C) inhibition. Nature. 2021;599(7886):679-83.

10. Misale S, Fatherree JP, Cortez E, Li C, Bilton S, Timonina D, et al. KRAS G12C NSCLC Models Are Sensitive to Direct Targeting of KRAS in Combination with PI3K Inhibition. Clin Cancer Res. 2019;25(2):796–807.

11. Amodio V, Yaeger R, Arcella P, Cancelliere C, Lamba S, Lorenzato A, et al. EGFR Blockade Reverts Resistance to KRAS(G12C) Inhibition in Colorectal Cancer. Cancer Discov. 2020;10(8):1129–39.

12. Feng J, Hu Z, Xia X, Liu X, Lian Z, Wang H, et al. Feedback activation of EGFR/wild- type RAS signaling axis limits KRAS(G12D) inhibitor efficacy in KRAS(G12D)-mutated colorectal cancer. Oncogene. 2023;42(20):1620–33.

13. Ryan MB, Coker O, Sorokin A, Fella K, Barnes H, Wong E, et al. KRAS(G12C)- independent feedback activation of wild-type RAS constrains KRAS(G12C) inhibitor efficacy. Cell Rep. 2022;39(12):110993.

14. Smith AE, Chan S, Wang Z, McCloskey A, Reilly Q, Wang JZ, et al. Tipifarnib Potentiates the Antitumor Effects of PI3Kalpha Inhibition in PIK3CA- and HRAS-Dysregulated HNSCC via Convergent Inhibition of mTOR Activity. Cancer Res. 2023;83(19):3252–63.

15. Adjei AA, Davis JN, Erlichman C, Svingen PA, Kaufmann SH. Comparison of Potential Markers of Farnesyltransferase Inhibition. Clinical Cancer Research. 2000;6(2318):2318–25.

16. Ho AL, Brana I, Haddad R, Bauman J, Bible K, Oosting S, et al. Tipifarnib in Head and Neck Squamous Cell Carcinoma With HRAS Mutations. Journal of Clinical Oncology. 2021;39(17):1856–64.

17. Baranyi M, Molnar E, Hegedus L, Gabriel Z, Petenyi FG, Bordas F, et al. Farnesyl- transferase inhibitors show synergistic anticancer effects in combination with novel KRAS-G12C inhibitors. Br J Cancer. 2024;130(6):1059–72.

18. Holderfield M, Lee BJ, Jiang J, Tomlinson A, Seamon KJ, Mira A, et al. Concurrent inhibition of oncogenic and wild-type RAS-GTP for cancer therapy. Nature. 2024;629(8013):919- 26.

19. Solanki HS, Welsh EA, Fang B, Izumi V, Darville L, Stone B, et al. Cell Type-specific Adaptive Signaling Responses to KRAS(G12C) Inhibition. Clin Cancer Res. 2021;27(9):2533–48.

20. Tanaka N, Lin JJ, Li C, Ryan MB, Zhang J, Kiedrowski LA, et al. Clinical Acquired Resistance to KRAS(G12C) Inhibition through a Novel KRAS Switch-II Pocket Mutation and Polyclonal Alterations Converging on RAS-MAPK Reactivation. Cancer Discov. 2021;11(8):1913–22.

21. Ryan MB, Fece de la Cruz F, Phat S, Myers DT, Wong E, Shahzade HA, et al. Vertical Pathway Inhibition Overcomes Adaptive Feedback Resistance to KRAS(G12C) Inhibition. Clin Cancer Res. 2020;26(7):1633–43.

22. Thatikonda V, Lyu H, Jurado S, Kostyrko K, Bristow CA, Albrecht C, et al. Co-targeting SOS1 enhances the antitumor effects of KRAS(G12C) inhibitors by addressing intrinsic and acquired resistance. Nat Cancer. 2024;5(9):1352–70.

23. Araujo HA, Pechuan-Jorge X, Zhou T, Do MT, Hu X, Rojas Alvarez FR, et al. Mechanisms of Response and Tolerance to Active RAS Inhibition in KRAS-Mutant Non–Small Cell Lung Cancer. Cancer Discovery. 2024:OF1-OF26.

24. Jänne PA, Bigot F, Papadopoulos K, Eberst L, Sommerhalder D, Lebellec L, et al. Abstract PR014: Preliminary safety and anti-tumor activity of RMC-6291, a first-in-class, tri- complex KRASG12C(ON) inhibitor, in patients with or without prior KRASG12C(OFF) inhibitor treatment. Molecular Cancer Therapeutics. 2023;22(12_Supplement):PR014-PR.

25. Gao H, Korn JM, Ferretti S, Monahan JE, Wang Y, Singh M, et al. High-throughput screening using patient-derived tumor xenografts to predict clinical trial drug response. Nat Med. 2015;21(11):1318–25.

